# Detecting genetically variant peptides in non-human samples

**DOI:** 10.1101/2024.08.22.609263

**Authors:** Fanny Chu, Andy Lin

**Affiliations:** Chemical and Biological Signatures, Pacific Northwest National Laboratory, Seattle, Washington 98109, United States

**Keywords:** forensic proteomics, liquid chromatography-tandem mass spectrometry, genetically variant peptides, contaminants

## Abstract

During proteomics data analysis, experimental spectra are searched against a user-defined protein database consisting of proteins that are reasonably expected to be present in the sample. Typically, this database contains the proteome of the organism under study concatenated with expected contaminants such as trypsin and human keratins. However, there are additional contaminants that are not commonly added to the database. In this study, we describe a new set of protein contaminants and provide evidence that they are detectable in mass spectrometry-based proteomics data. Specifically, we provide evidence that human genetically variant peptides (GVPs) can be detected in non-human samples. GVPs are peptides that contain single amino acid polymorphisms that result from non-synonymous single nucleotide polymorphisms in protein coding regions of DNA. We reanalyzed previously collected non-human data-dependent acquisition (DDA) and data-independent acquisition (DIA) datasets and detected between 0 and 135 GVPs per dataset. In addition, we show that GVPs are unlikely to originate from non-human sources and that a subset of eight GVPs are commonly detected across datasets.

## 1 Introduction

Proteomics researchers typically search experimentally-collected liquid chromatography-tandem mass spectrometry data against a user-defined protein database of expected sequences that aligns with the study design, e.g., proteome of the organism under study. This user-defined database usually contains the reference proteome of the sample being analyzed. For example, spectra generated from a human cell line would be searched against the human proteome. However, this approach may not account for the unintentional traces and their effects on analyte detection.

In addition to the reference proteome, the protein database is often augmented with contaminant proteins to account for proteins that are artificially-introduced into the sample during sample preparation. The widely known presence of contaminant proteins has spurred the development of standardized contaminant databases such as the CRAPome.^1^ More recently, work has been conducted to create universal contaminant libraries for both data-dependent acquisition (DDA) and data-independent acquisition (DIA) mass spectrometry data^2^ as well as affinity purification mass spectrometry.^3^ We note that these databases also contain contaminant sequences, such as trypsin and culture media peptides (e.g. bovine serum albumin), that are intentionally introduced during the sample preparation process but are irrelevant to the experimental question.

Even with the development of contaminant database, there are contaminants that are not commonly represented in the database. For example, chemical contaminants such as polyethylene glycol (PEG) are often not included within these databases.^4–6^

In this work, we describe a new set of protein contaminants and provide evidence that they are detectable in mass spectrometry-based proteomics data. Specifically, we provide evidence that mass spectrometry can detect contaminant human genetically variant peptides (GVPs) in non-human samples. GVPs are peptides that contain single amino acid polymorphisms that result from non-synonymous SNPs in protein coding regions of DNA. To date, all GVP detection has been performed on human samples, and to our knowledge, no one has investigated the feasibility of detecting human contaminant GVPs in non-human proteomics data.

Genetically variant peptide detection has been successfully performed on a variety of human samples. This approach was first proposed for use on human scalp hair in 2016.^7^ Since then, GVPs have been detected in hair obtained from different body locations,^8^ hair that had been damaged by an explosive event,^9^ and from single hairs.^10^ In addition, GVPs have been detected from other non-hair samples such as bone,^11^ plasma,^12^ and residues left from touch samples.^13, 14^ Furthermore, GVP detection has been successfully performed on data collected using different types of acquisition schemes such as data-dependent acquisition (DDA), data-independent acquisition (DIA), and parallel reaction monitoring (PRM).^15^

Traditionally, DIA data are searched against an experiment-specific spectral library that has created from previously collected data.^16, 17^ However, building experiment-specific spectral libraries is a time- and resource-intensive activity, and may not be feasible for low-quantity precious samples. As an alternative, recent work has proposed building predicted spectral libraries, where spectral prediction relies on machine learning models, such as Prosit^18^ and MS2Pip,^19^ that can predict fragment spectrum from peptide sequences.^20, 21^ However, to our knowledge, predicted spectral libraries have not been previously used for the detection of GVPs.

We demonstrate, by reanalyzing publicly-available non-human proteomics data, that mass spectrometry-based proteomics technology has the capability to detect GVPs in non-human samples. In addition, we provide evidence that GVPs can be detected in data acquired by DDA or DIA. Finally, we show that predicted spectral libraries can be used for contaminant GVP detection in DIA data.

## 2 Methods

Figure 1 provides a visual of the GVP detection workflow further described in the below sections. Spectra were annotated and visualized using spectrum utils.^22^

**Figure 1:**
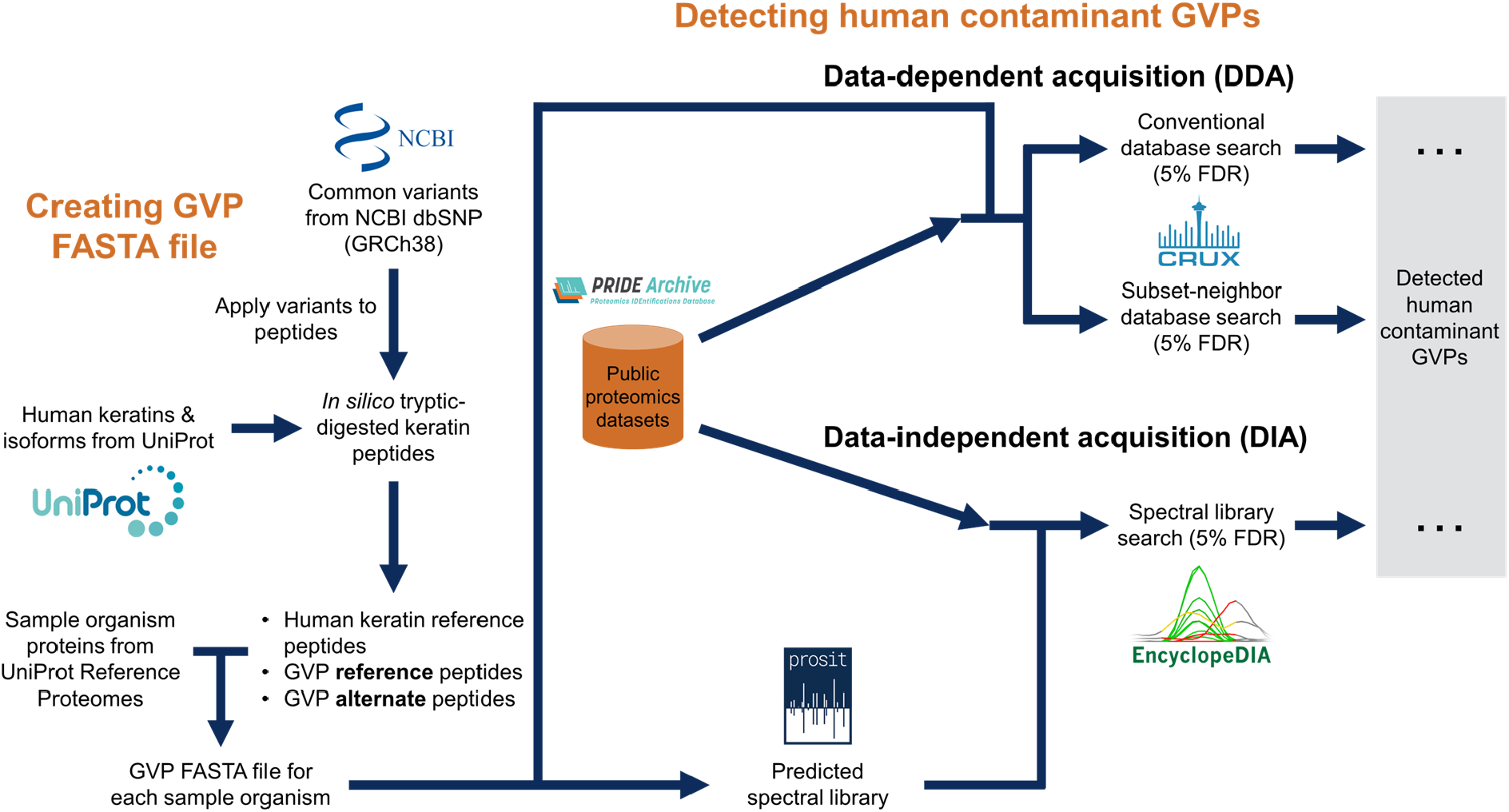
GVP detection workflow. Workflow for selection of PRIDE projects, GVP FASTA creation, and GVP detection.

### 2.1 Mass spectrometry datasets

Mass spectrometry runs from 18 different PRIDE projects (Supplementary Table S1 and S2)^23–40^ were downloaded from PRIDE.^41^ Of these 18 datasets, 12 were acquired by data-dependent acquisition (DDA) and six were acquired by data-independent acquisition (DIA). Within the DDA datasets, six datasets were acquired from *Escherichia coli* samples while the remaining six datasets were generated from yeast samples. For the DIA datasets, two were generated from *E. coli* samples while the other four were acquired on mouse samples.

### 2.2 Generating GVP fasta files

To search against mass spectrometry datafiles for peptide identification, FASTA files containing protein sequences from the sample organism and human contaminant genetically variant peptide sequences were constructed per sample organism. Protein sequences for sample organisms were obtained from UniProt.^42^ Specifically, the *E. coli*, yeast, and mouse proteomes were downloaded in May 2021, October 2021, and May 2021, respectively. Human contaminant genetically variant peptide sequences of interest were generated from a target list of common missense SNPs corresponding to human keratin-related proteins that are reported in the dbSNP database. Common missense SNPs are those SNPs that effect non-synonymous amino acid mutations and have global minor allele frequencies of at least 1% as reported by the Genome Aggregation Database (gnomAD).^43^ SNP consequences were then applied to all human keratin-related isoforms (GRCh38 build and obtained from Biomart) and mutated protein sequences were then subjected to *in silico* tryptic digest, to yield reference (unmutated) and alternate (mutated) GVP sequences. Human keratin reference peptides, which are human keratin sequences that do not contain SNPs, were also included in the FASTA file. SNPs that effect the Iso*→* Leu mutation, which does not result in a mass shift owing to the isobaric amino acids, were removed from consideration. Multi-allelic SNPs, which are mutations with more than two variants at a single location, were ignored, such that only the two most frequent variants at each location were considered (i.e., the major and minor alleles, also the reference and alternate alleles).

### 2.3 GVP identification from data-dependent acquisition data

Peptides were detected within DDA data via a database search using the Tide search engine^44^ implemented in Crux.^45, 46^ The precursor mass tolerance for each dataset was estimated using Param-Medic^47^ (Table S1) and was used to determine whether MS2 scans were acquired at high- or low-resolution. Runs that were collected with low-resolution MS2 scans were analyzed using the exact p-value score function,^48^ while runs that were collected with high-resolution MS2 scans were analyzed using the combined p-value score function.^49^ This dual score function approach was performed since exact p-value was developed for low-resolution data while combined p-value was developed for high-resolution data. For the searches using the exact p-value, we set `--score-function=exact-p-value`, `--exact-p-value=T`, and `--mz-bin-width=1.005079`. For the combined p-value searches, we set `--score-function=both`, `--exact-p-value=T`, and `--mz-bin-width=1.005079`. For all of these searches, all other parameters were set to their default values, except that `--top-match=1` and `--concat=T`. In addition, we allowed up to two methionine oxidation post-translational modifications per detected peptide.

We employed two different search strategies to analyze the data acquired by DDA. Specifically, we used the previously described subset-neighbor-search (SNS) and search-then-select methods.^50^ SNS was developed as a search strategy for when only a subset of peptides present in the sample are relevant. Briefly, spectra are searched against a database of relevant and neighbor peptides. A neighbor peptide is an irrelevant peptide that has similar precursor mass and fragmentation spectrum as a relevant peptide. In this study, we denote GVPs and keratin sequences that do not contain SNPs as relevant and any other peptide in the sample (such as species the sample originated from) as irrelevant. We use the same definition of a neighbor peptide as originally described, and we set *t*_*m*_ = 0.25 and *t*_*i*_ to be twice the precursor mass tolerance used in the associated database search.^50^ For high- and low-resolution searches, we used a bin size of 0.02 and 1.0005079 Da, respectively. Following the search, peptide-spectrum matches (PSMs) that involve neighbor peptides are removed and then the false discovery rate (FDR) is estimated on the remaining PSMs.

In addition to SNS, we also employed the search-then-select search strategy. This strategy is the standard approach and involves searching spectra against a database of peptides that are reasonably expected to be found in the sample. For this study, spectra were searched against a database that consisted of GVPs, the species proteome of the sample being analyzed, and human keratins. After the search, FDR was estimated on the resulting PSMs using the target-decoy competition strategy.^51^ Next, PSMs that do not involve a relevant peptides are removed, resulting in a final set of confident, relevant PSMs. While this strategy is found commonly throughout the literature, this method unfortunately does not guarantee proper control of the FDR on the subset of relevant PSMs.^50, 52, 53^ However, owing to its popularity, we nevertheless included this strategy.

For both search strategies, the FDR was jointly estimated over all runs within a PRIDE project using Crema.^54^ The FDR was estimated using target-decoy competition at both the PSM and peptide level, and filtered to a FDR threshold of 5%. For peptide-level FDR estimation, we employed the previously described “psm-and-peptide” methodology^55^ by setting `pep fdr type=psm-peptide`. The pairing file that is required for this method was generated by the tide-index tool within Crux by setting `--peptide-list=T`.

### 2.4 GVP identification from data-independent acquisition data

#### 2.4.1 GVP spectral library creation for data-independent acquisition data

Generalized spectral libraries for each sample organism were generated by using Prosit’s HCD model^18^ to predict peptide mass spectra from sequences in the FASTA files described in Section 2.2. Sequences in the FASTA files were tryptic-digested *in silico* in EncyclopeDIA^56^ (version 1.2) assuming the following generalized parameters: precursor charge state range 2 – 6, maximum of 2 missed cleavages, *m/z* range 345 – 1500, normalized collision energy 30, default charge state 3, and fragmentation method HCD. As of this writing, Prosit only considers a fixed carbamidomethylation modification.

#### 2.4.2 GVP identification from data-independent acquisition data using predicted spectral library

EncyclopeDIA^56^ (version 1.2) was used to search DIA mass spectrometry datasets against the GVP spectral library predicted by Prosit (Section 2.4.1). A target/decoy search approach was utilized, with precursor, fragment, and library mass error tolerances of 10 ppm. Identified precursors were then filtered to 5% FDR. Precursors were rolled up to peptide sequences and GVP sequences were further analyzed.

## 3 Results

We demonstrate the feasibility and ability to detect human contaminant GVPs in non-human samples, in both DDA and DIA datasets. The number of GVPs detected per PRIDE dataset, across all methods and datasets, ranged from 0–135 (Figure 2). In addition, we found there is a subset of eights GVPs that are commonly detected across many datasets. Finally, we determined that we can reasonably attribute human contaminant GVPs to human proteins and that a majority of these GVPs are not present in protein sequences of other organisms.

**Figure 2:**
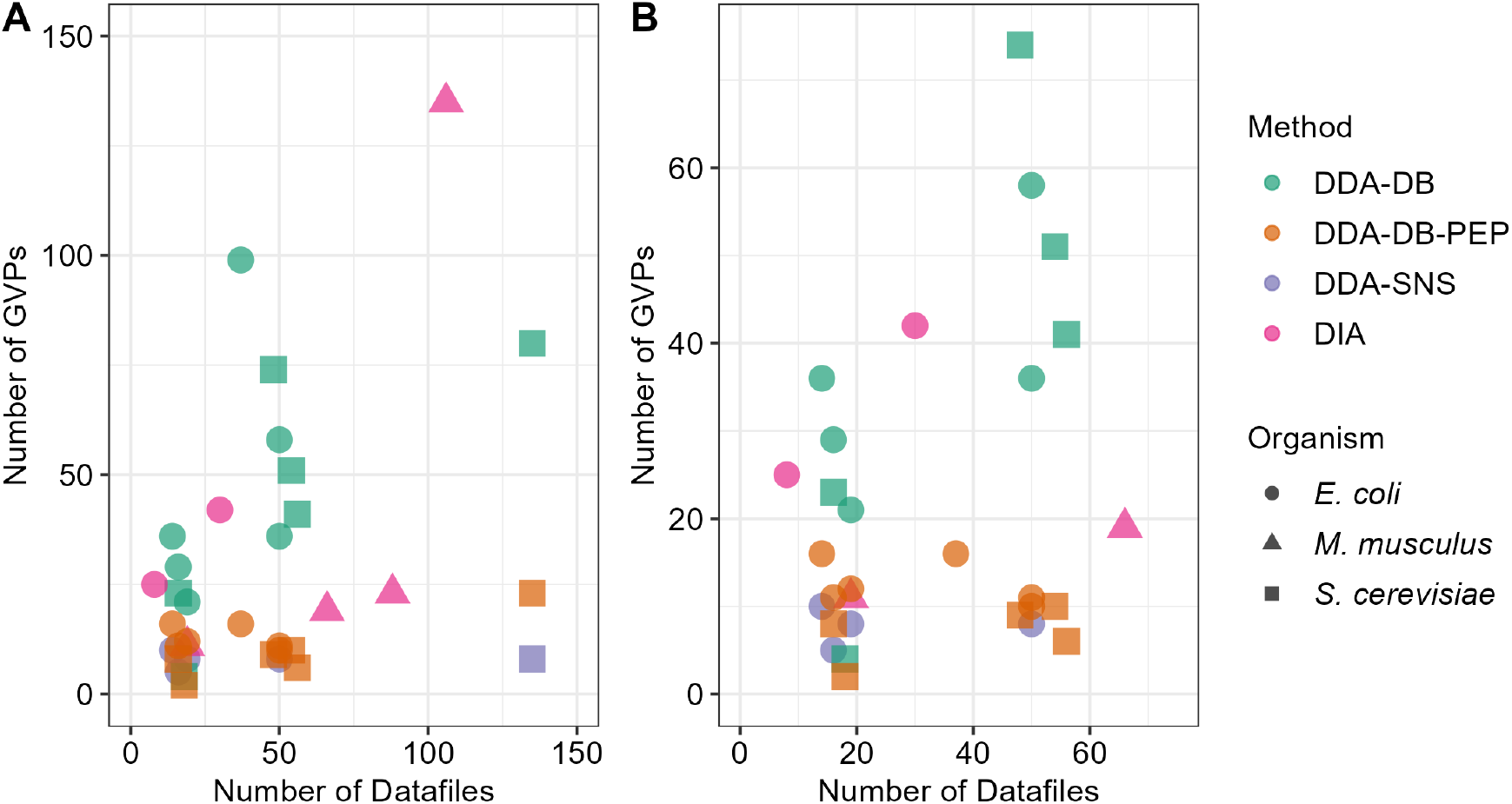
Number of GVP detection as a function of the number of datasets per PRIDE project. (A) A scatter plot of the number of GVP detections given the number of runs within each dataset. The shape indicates the species identity of the sample while the color represents the analysis method. (B) A subplot to focus on the lower-left quadrant of (A). Note that DDA-DB represents standard DDA database search analysis with PSM-level FDR control, DDA-DB-PEP represents the same with peptide-level FDR control, DDA-SNS represents subset-neighbor search with PSM-level FDR control, and DIA represents DIA analysis via spectral library search.

### 3.1 Human contaminant GVPs are detectable in DDA data of non-human organisms

First, we investigated the ability of mass spectrometry-based proteomics to detect human contaminant GVPs in non-human samples that have been acquired by DDA. We obtained and reanalyzed data from 12 projects on PRIDE,^41^ where each project contained between 14–135 runs (Table 1). Of the 12 PRIDE projects, six contained *E. coli* runs and six contained *S. cerevisiae* datasets (Table 1).

**Table 1:** Number of GVPs per project. A table of the number of detected GVPs and runs per PRIDE project. For data collected by DDA, the false discovery rate was estimated using the search-then-select procedure and filtered to a 5% peptide-level FDR.

| data acquisition method | sample organism | PRIDE ID | # of runs | # of GVPs |
| --- | --- | --- | --- | --- |
| DDA | *E. coli* | PXD010014 | 37 | 16 |
|  | *E. coli* | PXD016001 | 50 | 10 |
|  | *E. coli* | PXD022965 | 50 | 11 |
|  | *E. coli* | PXD023729 | 14 | 16 |
|  | *E. coli* | PXD025088 | 16 | 11 |
|  | *E. coli* | PXD025926 | 19 | 12 |
|  | yeast | PXD009420 | 16 | 8 |
|  | yeast | PXD014236 | 54 | 10 |
|  | yeast | PXD015273 | 48 | 10 |
|  | yeast | PXD015536 | 135 | 23 |
|  | yeast | PXD015674 | 18 | 2 |
|  | yeast | PXD026806 | 56 | 6 |
| DIA | *E. coli* | PXD010126 | 8 | 25 |
|  | *E. coli* | PXD012611 | 30 | 42 |
|  | mouse | PXD006240 | 88 | 23 |
|  | mouse | PXD016668 | 19 | 11 |
|  | mouse | PXD019379 | 66 | 19 |
|  | mouse | PXD027559 | 106 | 135 |

We utilized two database search strategies to determine whether we could detect human contaminant GVPs in existing runs. Specifically, we applied both the conventional database search approach, which has been previously termed “search-then-select”, and “subset-neighbor-search” (SNS) approach.^50^ SNS was developed to address the case where a subset of the peptides in the sample are relevant to downstream analysis (i.e., PSMs corresponding to GVPs). We note that previous work has shown that the conventional approach is not guaranteed to control the FDR of the relevant subset of PSMs.^50^ While the standard approach is not guaranteed to control the FDR, we opted to include this approach due to its ubiquity.

In addition to two different database search approaches, we also investigated GVP detection when the FDR was controlled at either the psm-level or the peptide-level. We note that while PSM-level FDR is traditionally employed, recent work has shown that this approach may not properly the FDR and that peptide-level FDR should be used instead.^55, 57^

We found that were successfully able to detect GVPs in all 12 DDA datasets when we used the search-then-select procedure with a 5% peptide-level FDR (Table 1). We detected an average of 11.25*±*5.10 (s.d.) GVPs (min=2; max=23) per project. Across all 12 datasets, a total of 76 different GVPs were detected. Out of these 76 GVPs, 17 were detected in both *E. coli* and yeast runs (Supplementary Figure S1). Within the set of 17, 12 originated from reference alleles. The vast majority of these GVPs belong to keratins (11 different keratins), many of which are cytoskeletal keratins. While we were able to detect GVPs using the search-then-select procedure, no GVPs were detected when the SNS procedure was performed with peptide-level FDR control.

In addition to peptide-level FDR control, we were able to successfully detect GVPs in all 12 DDA dataset when we used the search-then-select procedure with a 5% PSM-level FDR control. Specifically, we detected an average of 159.92*±*109.78 (s.d.) GVP PSMs (min=14; max=423) per project. In addition, when considering base unique sequences only, we detected an average of 45.17*±*27.05 (s.d.) unique GVP sequences (min=4; max=99) (Supplemental Table S3). Across all the runs, we detected a total of 285 different GVPs. A subset of 87 GVPs were found in runs from both species, and these GVPs originated from 44 different genes (Supplementary Figure S2). Additionally, a majority of the GVPs (35/44; 79.55%) detected across species were reference GVPs. Notably, many GVPs belong to filaggrin and other structural proteins, though other detected GVPs derive from cytoskeletal as well as cuticular keratins. Of the 27 keratins observed from profiling the common GVPs in DDA data, 10 are hair keratins (i.e., cuticular), 12 are epithelial keratins, and remainder are hair follicle-specific epithelial keratins.^58^ A majority of detected GVPs deriving from epithelial keratins and keratins known to be highly expressed in skin aligns with expectations of human contamination during sample handling, such as from shed skin cells. Finally, GVPs were only confidently detected in one yeast dataset and four *E. coli* datasets when the SNS protocol was used in conjunction with a 5% PSM-level FDR control (Supplemental Table S3).

Due to some of the statistical concerns with the search-then-select methodology, we opted to perform validation of these detections. Specifically, we manually validated the best and worst scoring detection for each of the 12 datasets (Supplemental Figure S4–S15). This validation step was performed when the FDR was controlled at PSM-level and peptide-level. After visualizing the annotated spectra, we accepted all 12 best scoring detections as being correct for both PSM-level and peptide-level FDR. On the other hand, we accepted three and four of the worst-scoring PSMs as being correct when controlling for PSM-level and peptide-level FDR, respectively. This indicates that, while not all detections may be correct, it is possible for mass spectrometry-based proteomics to detect contaminant GVPs. Note that the best scoring PSM-level detection and the peptide-level detection are the same detection.

### 3.2 Human contaminant GVPs can be detected in DIA data of non-human organisms

After determining that human contaminant GVPs can be detected in non-human samples acquired by DDA, we hypothesized that it would be also possible to detect human contaminant GVPs in non-human samples acquired by DIA. To test this, we reanalyzed six previously acquired datasets (four mouse and two *E. coli* projects) that were publicly available on PRIDE using a spectral library approach. Specifically, we searched the data using EncyclopeDIA against a Prosit-predicted spectral library.

We found that we were able to detect human contaminant GVPs in non-human samples that had been acquired by DIA. Of the six PRIDE projects examined, which contained between 4–106 runs per project, an average of 43 *±* 46 GVPs (s.d.) were detected per project (Table 1 and Figure 2).

Out of the total of 189 GVPs detected across all six DIA PRIDE projects, we found 37 GVPs that were present in both *E. coli* and mouse datasets (Figure 3). These 37 GVPs originated from 23 genes, and the vast majority of commonly detected GVPs belonged to the reference allele. However, in a few cases (i.e., KRT13, KRT23, and KRT76), both the reference and alternate allele of the same SNP were detected. Again, as expected, the majority of GVPs belong to cytoskeletal keratins, 12 cytoskeletal keratins represented and 3 cuticular keratins; a number of these keratins are known to be highly expressed in skin.

**Figure 3:**
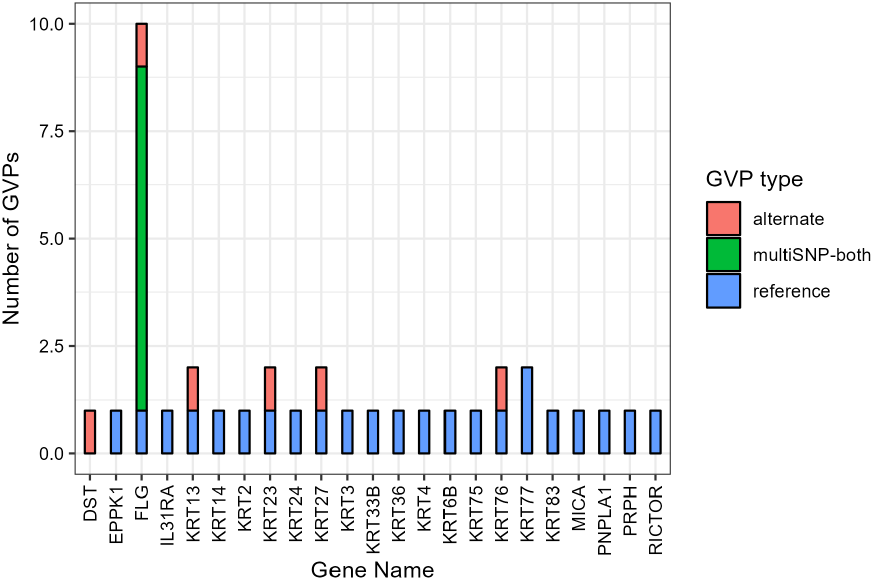
GVPs detected found in DIA data. The distribution of common GVPs detected in DIA data by gene name and GVP type. A common GVP is a GVP that is detected in at least one *E. coli* and mouse dataset. A GVP labeled as “multiSNP-both” indicates a tryptic peptide with multiple non-synonymous sites that where at least one site is a reference allele and one site is an alternate allele.

### 3.3 Determining frequently detected GVPs

After determining that GVPs can be detected in previously acquired DDA and DIA data, we set out to determine whether certain GVPs are frequently detected across datasets. In addition, we investigated whether any GVPs were detected across acquisition style and sample organism at a 5% FDR. For the DDA data, we considered confident detections obtained using the search-then-select approach with peptide-level FDR control.

Our results suggests that there are a subset of GVPs that are commonly detected across datasets. Specifically out of the 236 GVPs that we detected across 18 different projects, we identified eight GVPs that were detected in at least five different PRIDE projects (Figure 4 and Table 2). These eight common GVPs were detected in anywhere between five to 15 projects and were detected in samples collected by both DDA and DIA. Specifically, these common GVPs were present in one to 11 DDA projects (out of 12) and one to five DIA projects (out of six). In addition, we found that these GVPs were detected regardless of sample organism. Generally, we found that these common GVPs were detected in all three sample organisms that we analyzed (yeast, mouse, and *E. coli*). One exception was that AGLETAIADAEQR was not detected in *E. coli* runs collected by DDA or DIA. The other exception was that FLEQQNQVLETK was not detected in *E. coli* runs collected by DIA.

**Table 2:**
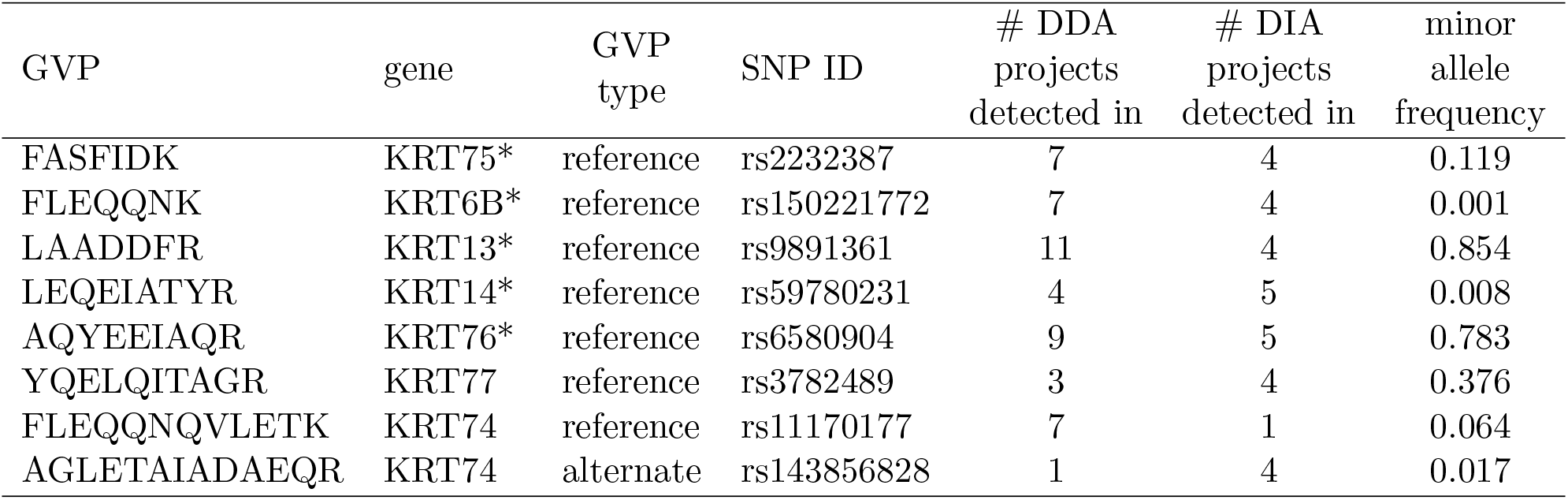
Common GVPs. A table of common GVPs where a common GVP was detected in at least five different PRIDE projects. There are a total of 12 DDA projects and 6 DIA projects. The major allele frequency is in the context of all human ancestry. *These sequences are present in reference sequence of other keratin genes (see Supplementary Table S4 for more details).

**Figure 4:**
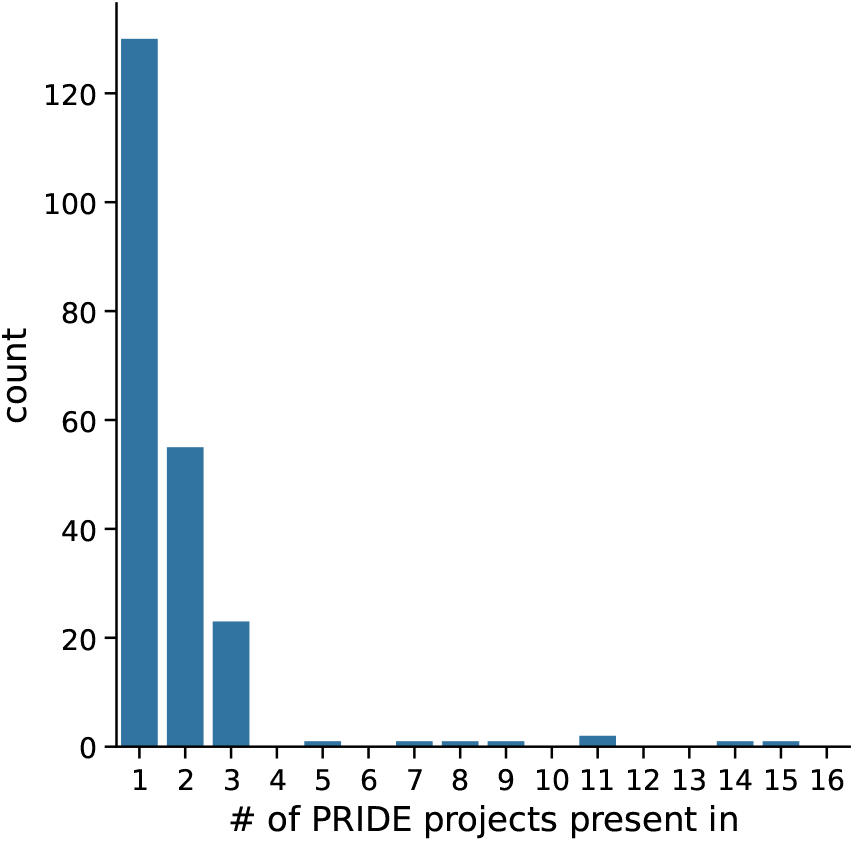
Identification of common GVPs. The number of PRIDE projects each GVP is detected in. Most GVPs are found in few datasets. However, there are eight GVPs that are commonly detected across the datasets analyzed.

Further investigation of these common GVPs found that they all originated from cytoskeletal keratins as opposed to cuticular keratins, the latter of which are highly expressed in hair (Table 2). In addition, we found that a large majority of these GVPs (7/8) originated from the reference allele. When using the global population, the detected common reference GVPs had minor allele frequencies ranging from 0.001–0.854 while the alternate GVP had a minor allele frequency of 0.017. In general, common GVPs had high allele frequencies, which matches our expectations that GVPs with high allele frequencies are more likely to be detected across projects. One exception was the GVP associated with an alternative allele, which had an allele frequency of 0.017. This is unexpected as this low allele frequency would indicate that it would be unlikely to be detected in a large number of projects. However, we note that this GVP was detected in the fewest number of projects out of the set of eight common GVPs. In addition, we note that this GVP was detected in four DIA projects and one DDA project. It is unclear why this alternate GVP was detectable in so many DIA datasets. However, we do not believe this peptide originates from unexpected non-human contamination as the sequence was not found in Uniprot. Future work could be pursued to determine if different acquisition strategies yield different set of detected GVPs.

### 3.4 Most human contaminant GVPs are not found in the proteomes of other organisms

We have shown evidence that it is possible to detect human contaminant GVPs in non-human samples. However, an alternative hypothesis to this finding is that these putative human contaminant GVPs could instead originate from other non-human protein contaminants. To investigate the possibility of detected human contaminant GVPs instead derive from non-human sources, we examined whether sequences are unique with respect to human and non-human proteomes.

To confirm the uniqueness of human contaminant GVPs compared to proteomes of other organisms, we used UniPept^59, 60^ to perform a lowest common ancestor (LCA) analysis on all theoretically possible reference and alternative human keratin-related GVPs. In an LCA analysis, all of the species that a peptide could originate from is collected. Next, the lowest common taxonomic rank of these species is determined and assigned to each sequence.

After determining the LCA of each GVP, we bin them into five different categories. Peptides assigned to the “*H. sapiens*” bin are unique to human proteins. In addition, peptides assigned to the “*H. sapiens* lineage” are found in proteins from species that have the same taxonomic lineage as human from the domain to the genus level. Furthermore, peptides assigned to “other lineage” are shared with proteins from organisms not in the *H. sapiens* taxonomic lineage. Finally, peptides assigned to “root” or “unmatched” are shared with proteins common to all organisms or not found in any proteins within the database, respectively.

If the alternative hypothesis is true, then the vast majority of the detected human contaminant GVPs should not be unique to the human proteome and thus, be found in the proteomes of other organisms outside of the human taxonomic lineage, that is, “other lineage”. But if these sets of human contaminant GVPs do indeed derive from human proteins, then they should be assigned to the “*H. sapiens*” or “*H. sapiens* lineage” bins.

We found that a vast majority of reference GVPs are either shared with proteins in the *H. sapiens* lineage or unique to human proteins (Figure 5). Specifically, 83 (23.9%) and 139 (40.0%) out of 347 reference GVPs were unique to humans or mapped to the *H. sapiens* lineage, respectively. On the other hand, a majority of alternate GVPs (289 out of 426) are unmatched to any sequence in Unipept (Figure 5). We theorize this may be due to UniPept only containing canonical protein sequences and are therefore more likely to contain reference instead of mutated peptides; mutated peptides are typically captured as an isoform to the canonical sequence.

**Figure 5:**
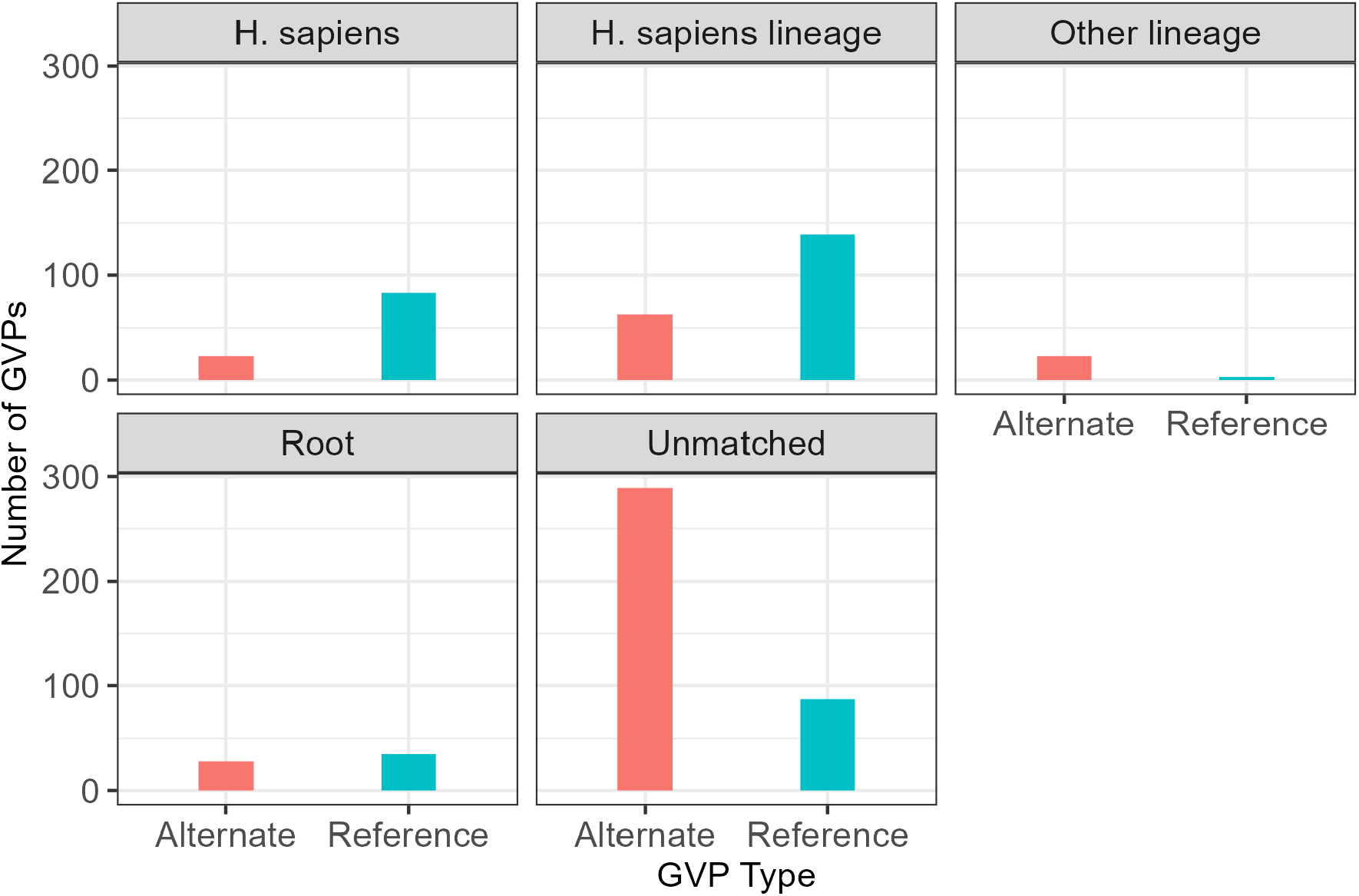
Lowest common ancestor analysis. A lowest common ancestor analysis of all theoretical reference and alternate human keratin-related GVPs using UniPept. A “Unmatched” GVP represents a peptide sequence not found in UniPept while a “*H. sapiens* lineage” GVP is a GVP that is only found in the taxonomy lineage of *H. sapiens*.

In addition, we found a few GVPs that mapped to the root or non-human taxonomic lineages. Specifically, 35 reference and 28 alternate GVPs mapped to the root, while 3 reference and 23 alternate GVPs mapped to non-human taxonomic lineages. It is unsurprising that alternate GVPs could map to non-human taxonomic lineages as Unipept likely only contains canonical protein sequences. However, investigation of the implications of this observed ambiguity on GVP detection and protein assignment is outside the scope of this study.

On the other hand, three reference human keratin-associated GVPs unexpectedly mapped to non-human taxonomic lineages. These three sequences (PTCCQNTCCR, AFSCISACGPR, STYYQPICYIFK) mapped to a taxonomic lineage of *Pan paniscus, Aotus nancymaae*, and *Caniformia*, respectively. Further investigation of these sequences determined that the first peptide has a proline at the beginning of the sequence while the next amino acid of the other two sequence are also a proline. Therefore, we hypothesize that these peptides map to non-human taxonomic lineages owing to differences in the application of the proline rule. Specifically, we did not use the proline rule when generating GVPs while UniPept employs the proline rule.^61^

Aligning with our expectations, we observed that reference GVPs typically mapped to a human taxonomic lineage and alternate GVPs are unmatched to any organism. This finding provides confidence that GVPs detected in the proteomics data of non-human organisms likely originate from human contamination rather than being derived from other non-human organism contaminants.

## 4 Discussion

In this work, we investigate the ability of mass spectrometry-based proteomics to detect human contaminants GVPs. We demonstrated that MS-based proteomics can detect these GVPs across a variety of projects and sample types. In addition, we showed that these GVPs are unlikely to originate from unexpected non-human sources and also determined which GVPs are commonly detected across datasets.

While we have shown that MS-based proteomics can detect human contaminant GVPs, this capability is still in its infancy and there are numerous potential avenues of effort to pursue. One potential avenue is to investigate whether putative GVPs detections could instead result from post-translational modifications. We observed 2 different GVP sequences, detected in two different mass spectrometry runs, in which the peptide may be ambiguously identified, as they exhibit single amino acid polymorphisms that coincide with the common post-translational modification deamidation. Given the scarce instances of these occurrences in the datasets we examined, it is unlikely to be a pervasive issue. Nevertheless, future work could investigate amino acid polymorphisms with a larger set of modifications.

In addition, we note that because our work reanalyzed existing data, we are unable to confirm our GVPs detections with genomic evidence. Future work could determine the sensitivity and specificity through analysis of ground truth sample. In addition, analysis of ground truth samples could confirm that GVPs originate from sample handlers.

Finally, this effort helps increase the proportion of spectra that are identified from a run. Despite decades of work, it is common for a large number of spectra within a run to be remain unidentified.^62^ These unidentified spectra are often referred to the “dark matter” of the proteome or the “dark proteome”. Work has been done to elucidate the dark proteome. For example, one approach has focused on doing so via identification of novel post-translational modifications.^63^ Our work suggests that continued investigation into contaminants may be a complementary method to advance elucidation of the dark proteome.

## 5 Acknowledgments

This research was supported by the Laboratory Directed Research and Development Program at Pacific Northwest National Laboratory, a multiprogram national laboratory operated by Battelle for the U.S. Department of Energy. Andy Lin is grateful for the support of the Linus Pauling Distinguished Postdoctoral Fellowship program. Fanny Chu is grateful for the support of the Open Call Initiative. We would also like to thank PNNL Research Computing for access to computational resources. Pacific Northwest National Laboratory is a multiprogram national laboratory operatedn by Battelle Memorial Institute for the United States Department of Energy under contract DE-AC06-76RLO.

## 6 Supporting Information

- Supplemental File S1: PDF containing supplemental tables and figures.

## For Table of Contents Only

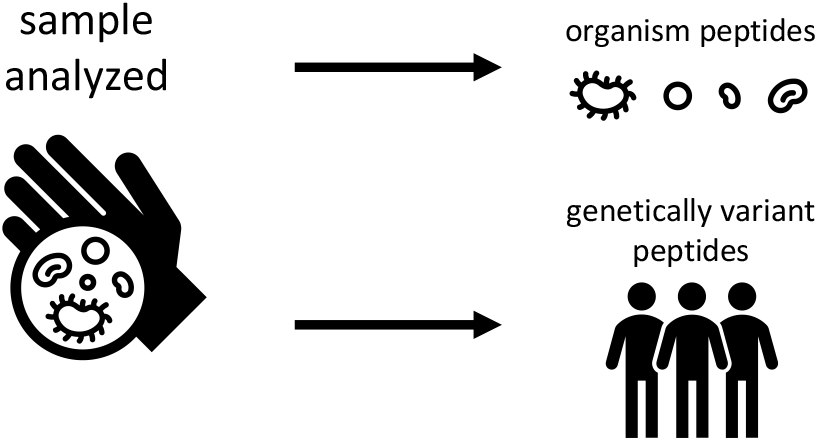

