## Supplemental File for "Detecting genetically variant peptides in non-human samples"

August 9, 2024

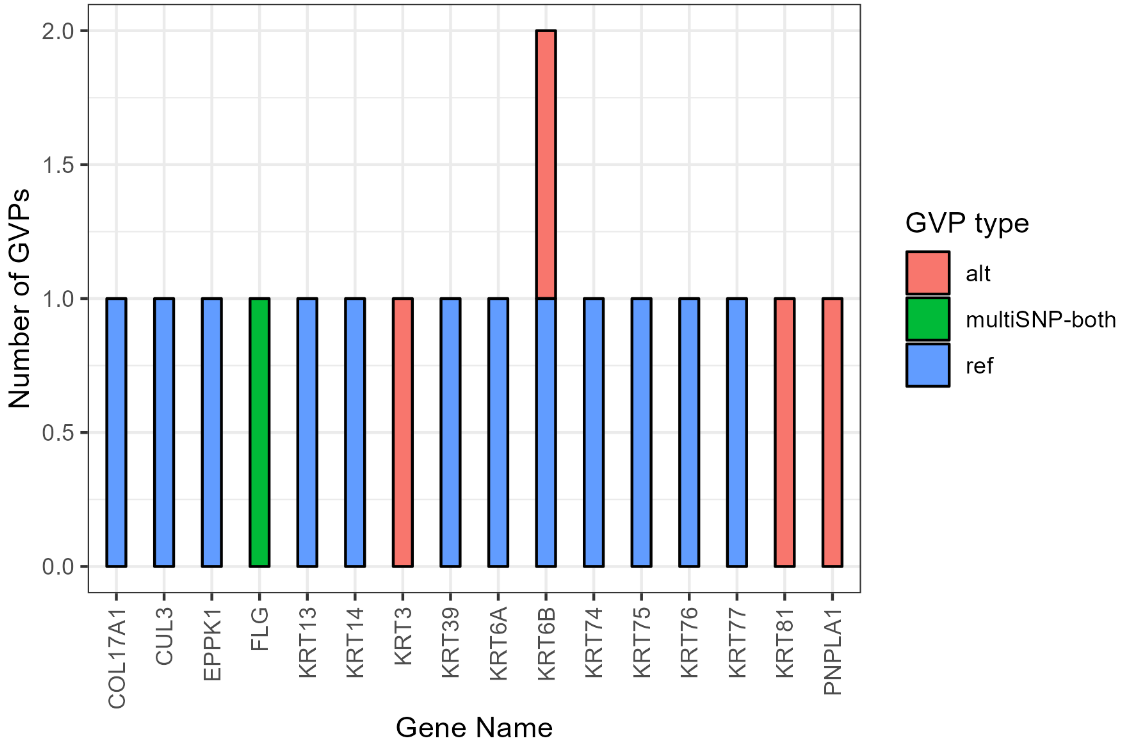


Figure S1: **Common GVPs in DDA data with peptide-level FDR control**. The genes where commonly detected GVPs from DDA are located in. We define a commonly detected GVP as one that is found in both *E. coli* and yeast datasets. Peptide detections were filtered to a 5% peptide-level FDR using the search-then-select approach.

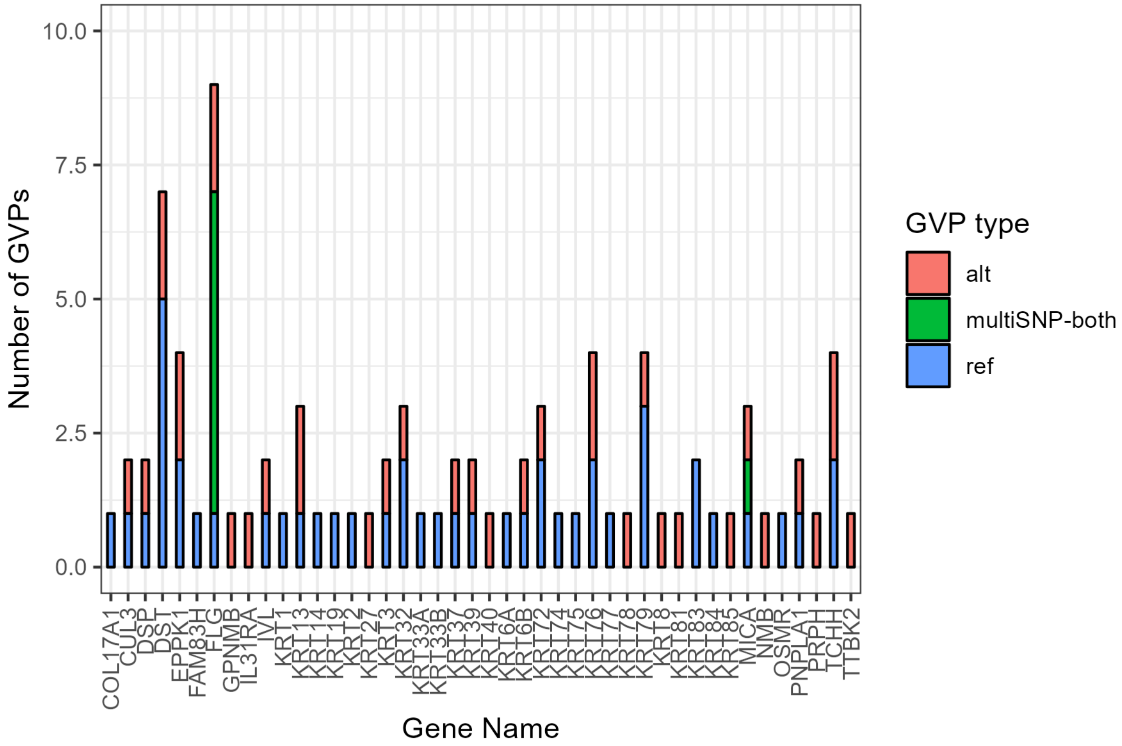


Figure S2: **Common GVPs in DDA data with PSM-level FDR control** The genes where commonly detected GVPs from DDA are located in. We define a commonly detected GVP as one that is found in both *E. coli* and yeast datasets. Peptide detections were filtered to a 5% PSM-level FDR using the search-then-select approach.

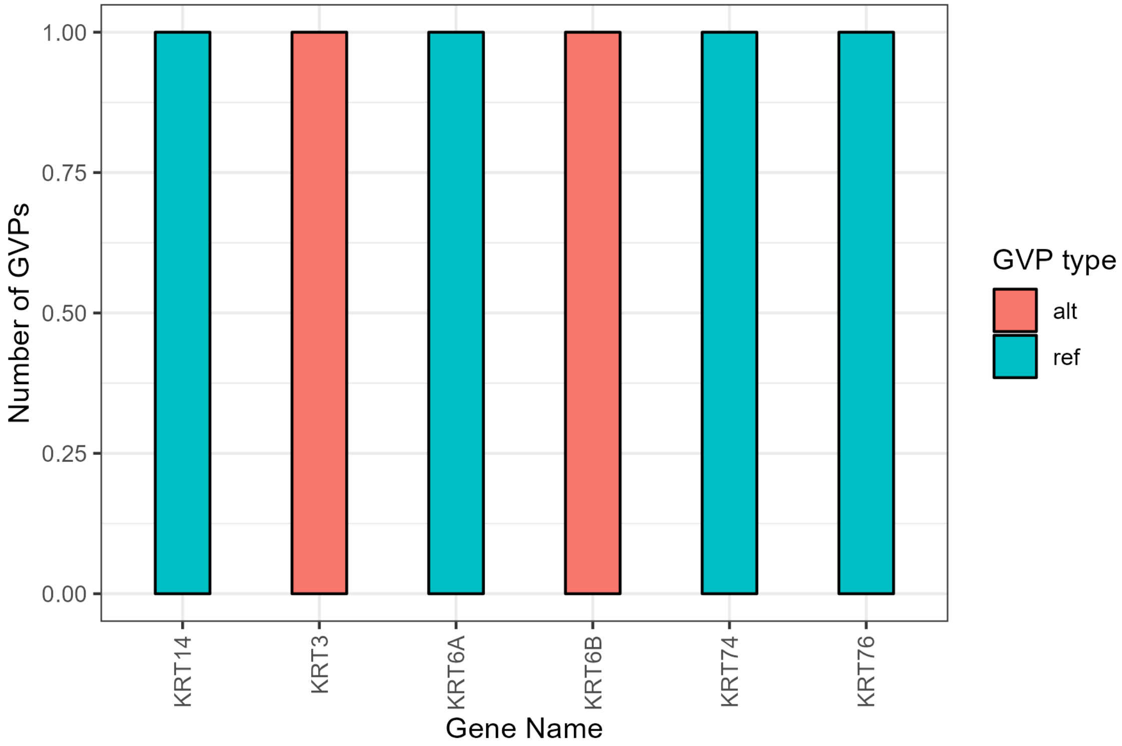


Figure S3: **Common GVPs in DDA data with PSM-level FDR control** The genes where commonly detected GVPs from DDA are located in. We define a commonly detected GVP as one that is found in both *E. coli* and yeast datasets. Peptide detections were filtered to a 5% PSM-level FDR using the subset-neighbor-search approach.

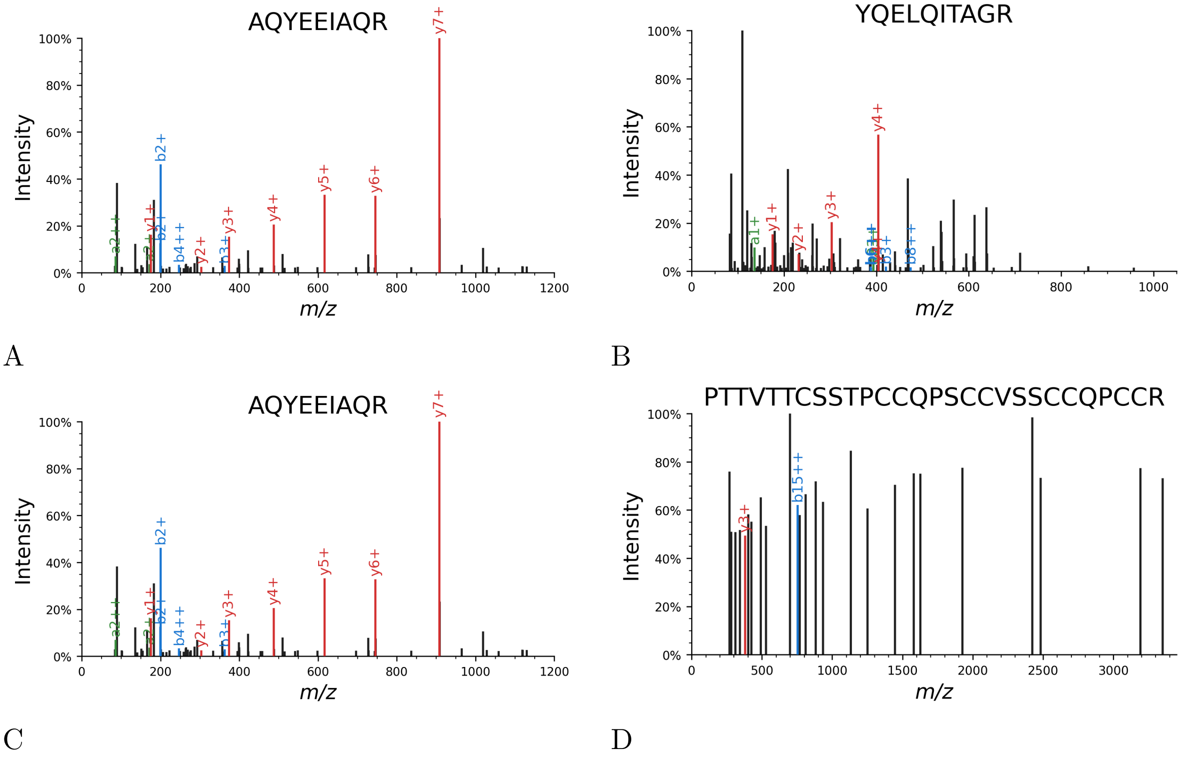


Figure S4: **Annotated spectra for project PXD009420.** Annotated spectra of the best (A) and worst (B) scoring confident detection when controlling the PSM-level FDR using the search-then-select protocol. We also the annotated spectra of the best (C) and worst (D) confident detection when controlling the peptide-level FDR.

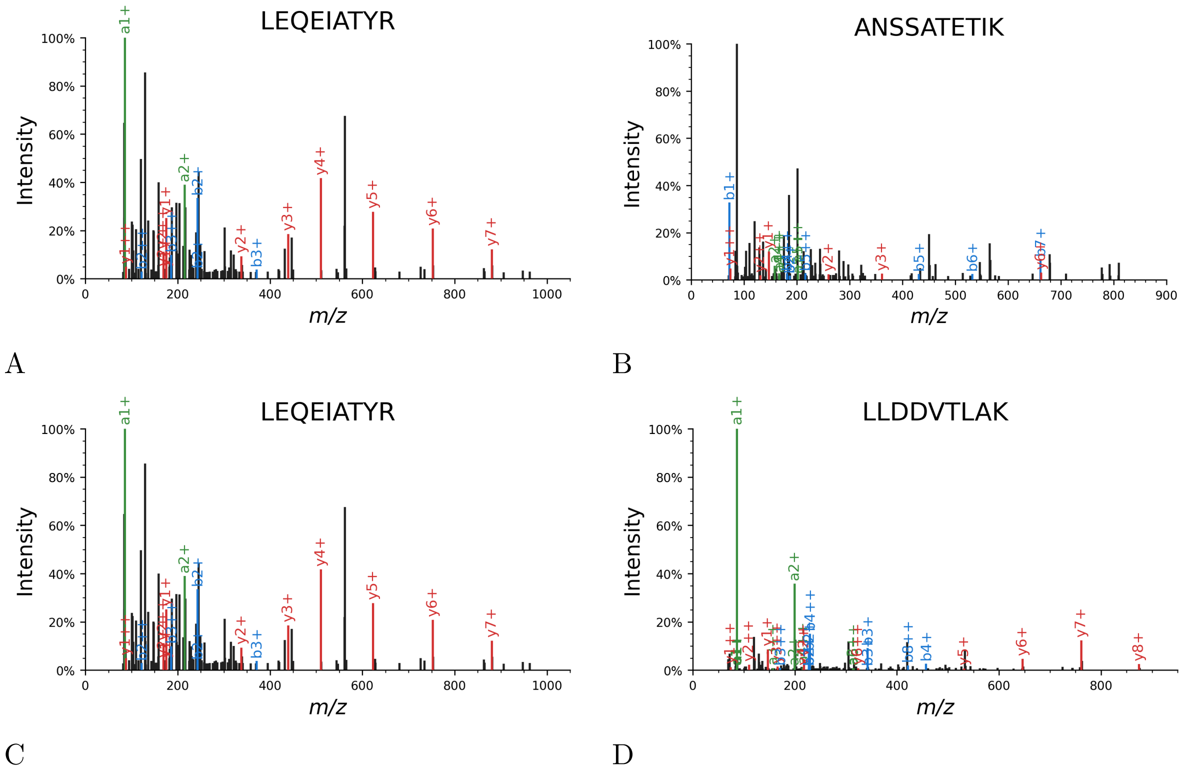


Figure S5: **Annotated spectra for project PXD014236.** Annotated spectra of the best (A) and worst (B) scoring confident detection when controlling the PSM-level FDR using the search-then-select protocol. We also the annotated spectra of the best (C) and worst (D) confident detection when controlling the peptide-level FDR.

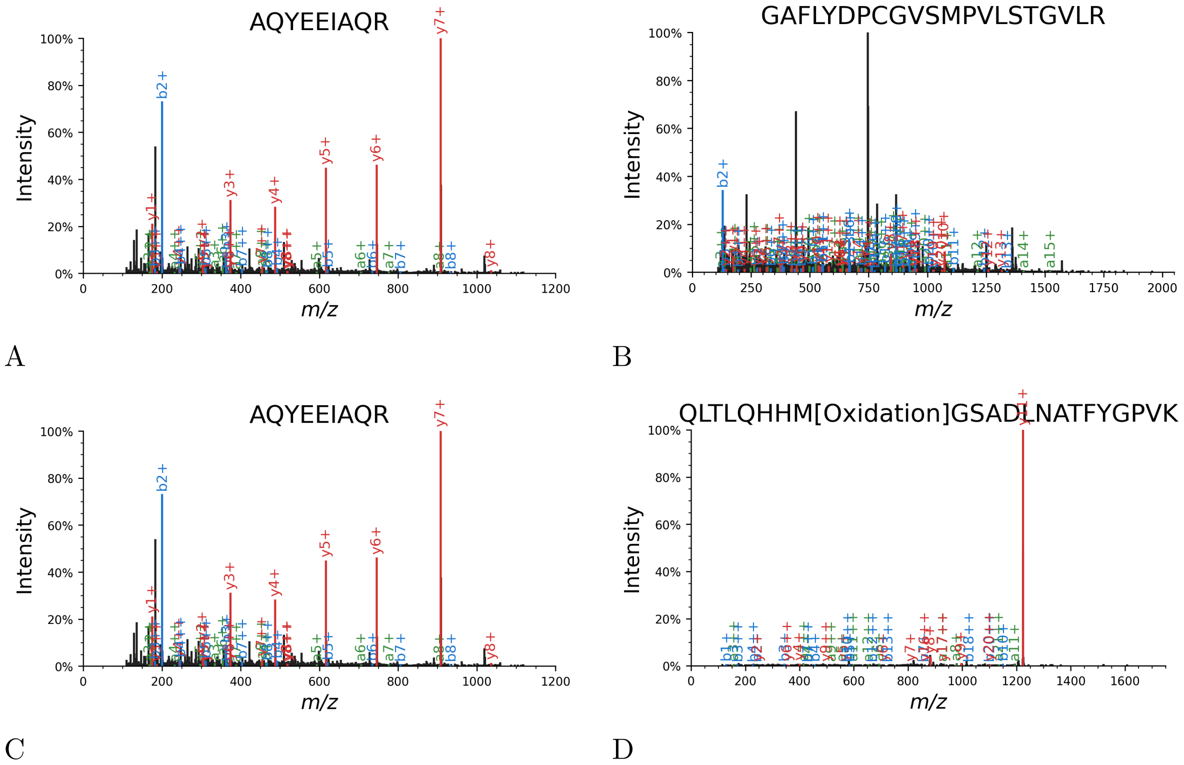


Figure S6: **Annotated spectra for project PXD015273.** Annotated spectra of the best (A) and worst (B) scoring confident detection when controlling the PSM-level FDR using the search-then-select protocol. We also the annotated spectra of the best (C) and worst (D) confident detection when controlling the peptide-level FDR.

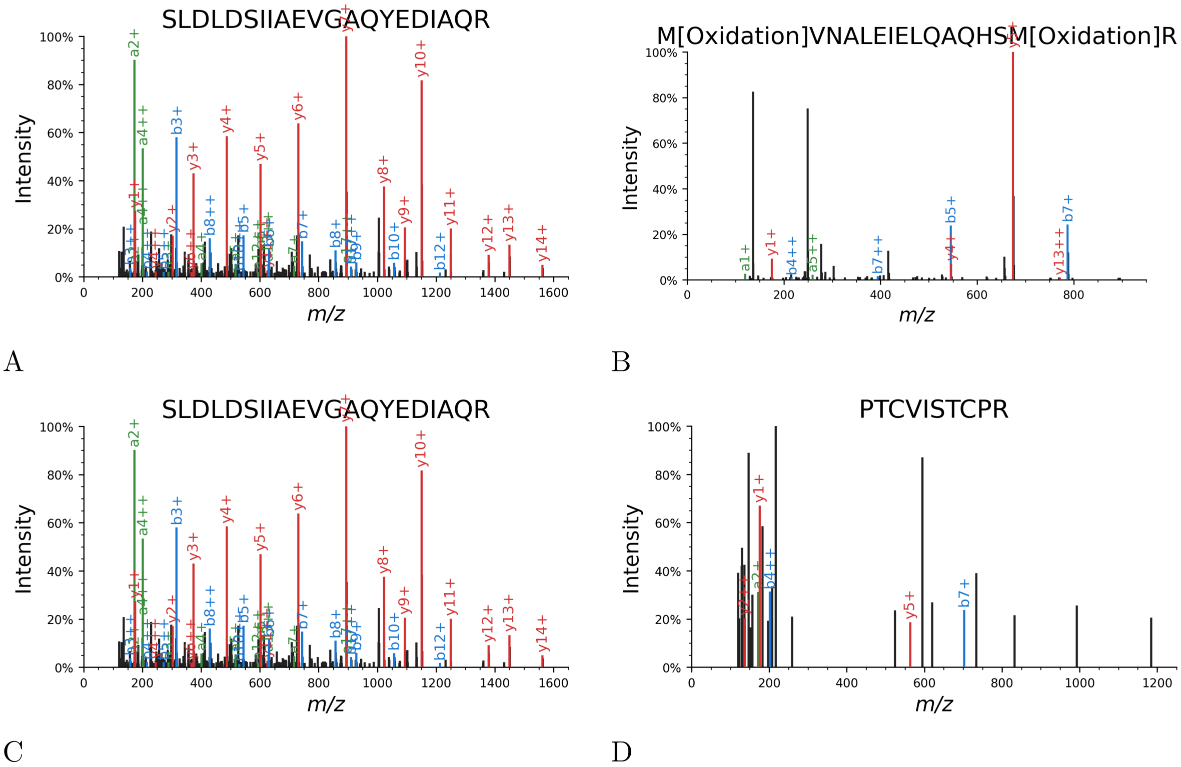


Figure S7: **Annotated spectra for project PXD015536.** Annotated spectra of the best (A) and worst (B) scoring confident detection when controlling the PSM-level FDR using the search-then-select protocol. We also the annotated spectra of the best (C) and worst (D) confident detection when controlling the peptide-level FDR.

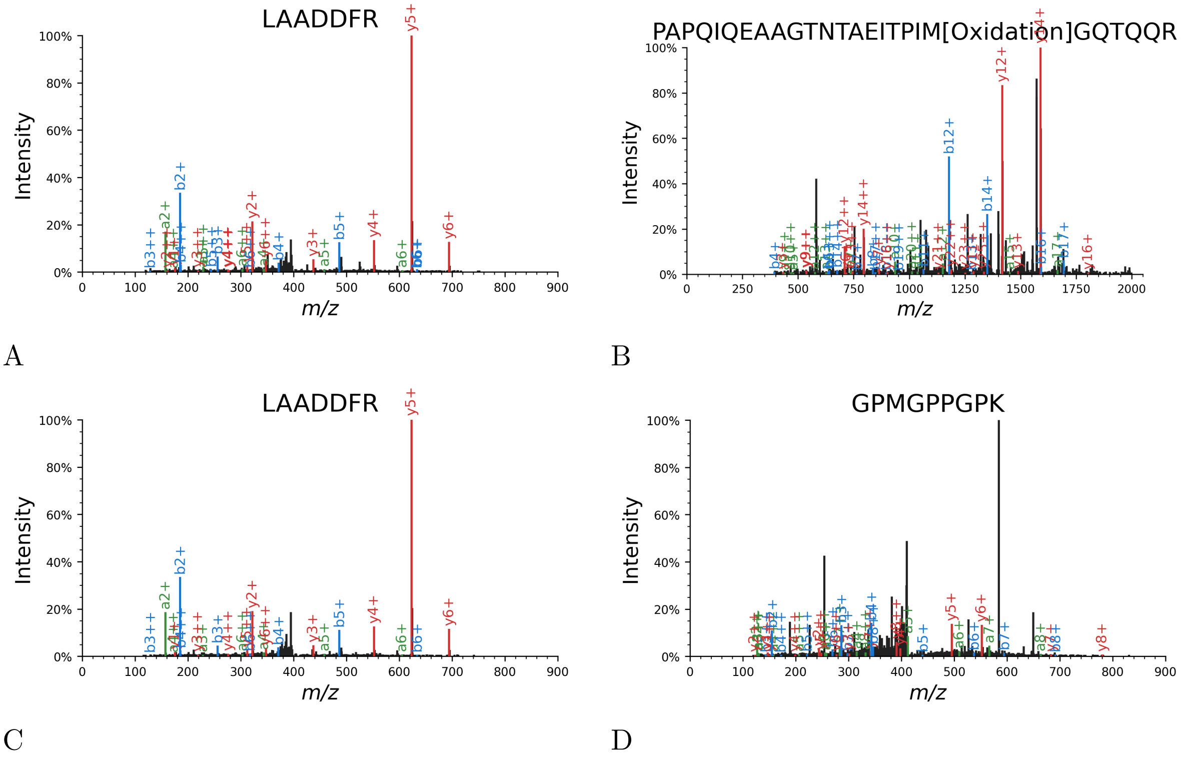


Figure S8: **Annotated spectra for project PXD015674.** Annotated spectra of the best (A) and worst (B) scoring confident detection when controlling the PSM-level FDR using the search-then-select protocol. We also the annotated spectra of the best (C) and worst (D) confident detection when controlling the peptide-level FDR.

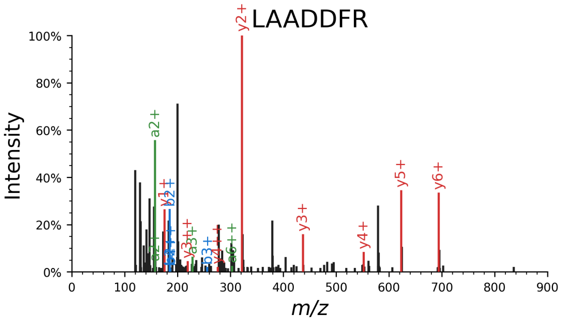


A

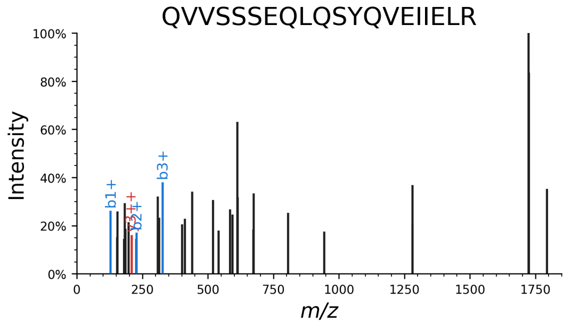


B

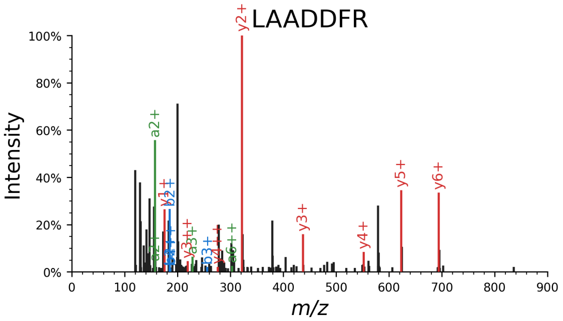


C

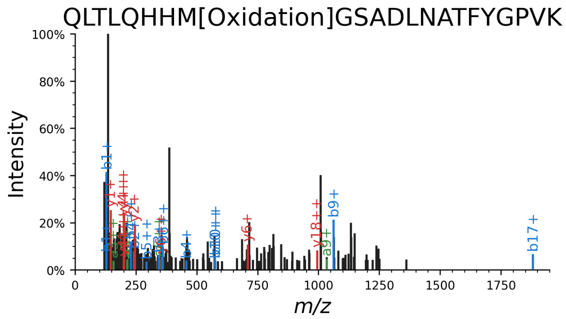


D

Figure S9: **Annotated spectra for project PXD026806.** Annotated spectra of the best (A) and worst (B) scoring confident detection when controlling the PSM-level FDR using the search-then-select protocol. We also the annotated spectra of the best (C) and worst (D) confident detection when controlling the peptide-level FDR.

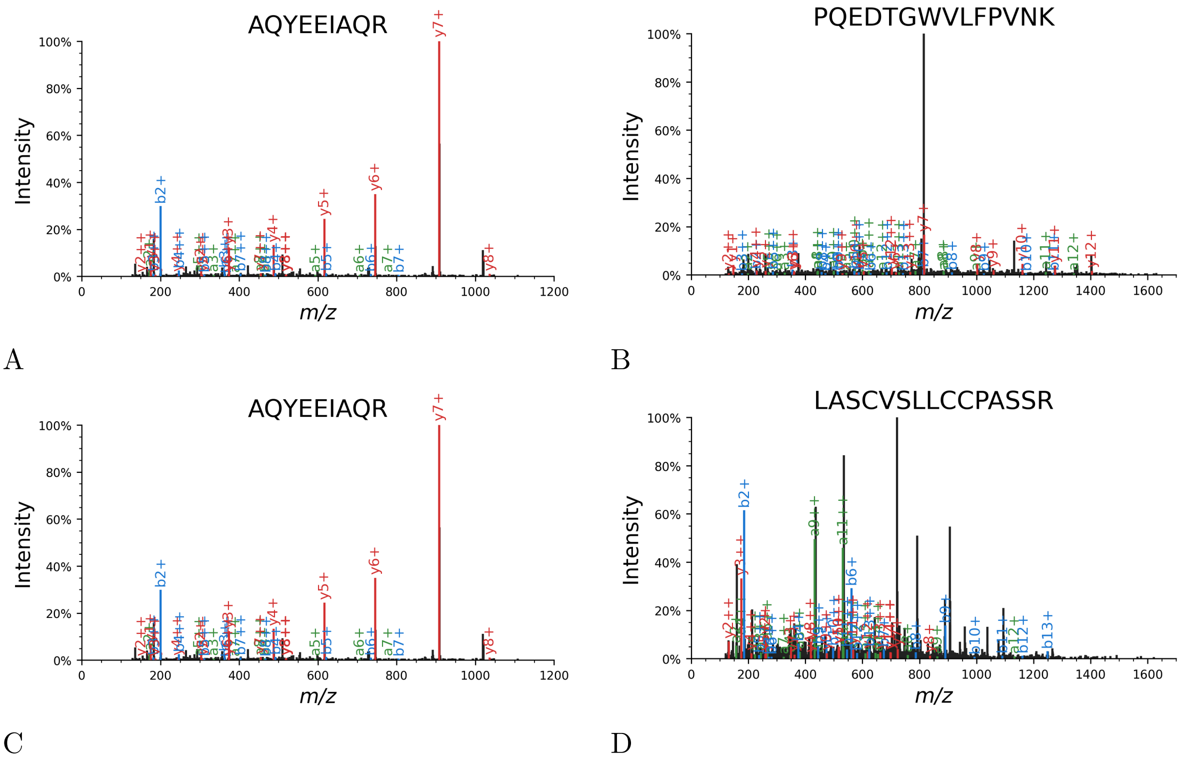


Figure S10: **Annotated spectra for project PXD010014.** Annotated spectra of the best (A) and worst (B) scoring confident detection when controlling the PSM-level FDR using the search-then-select protocol. We also the annotated spectra of the best (C) and worst (D) confident detection when controlling the peptide-level FDR.

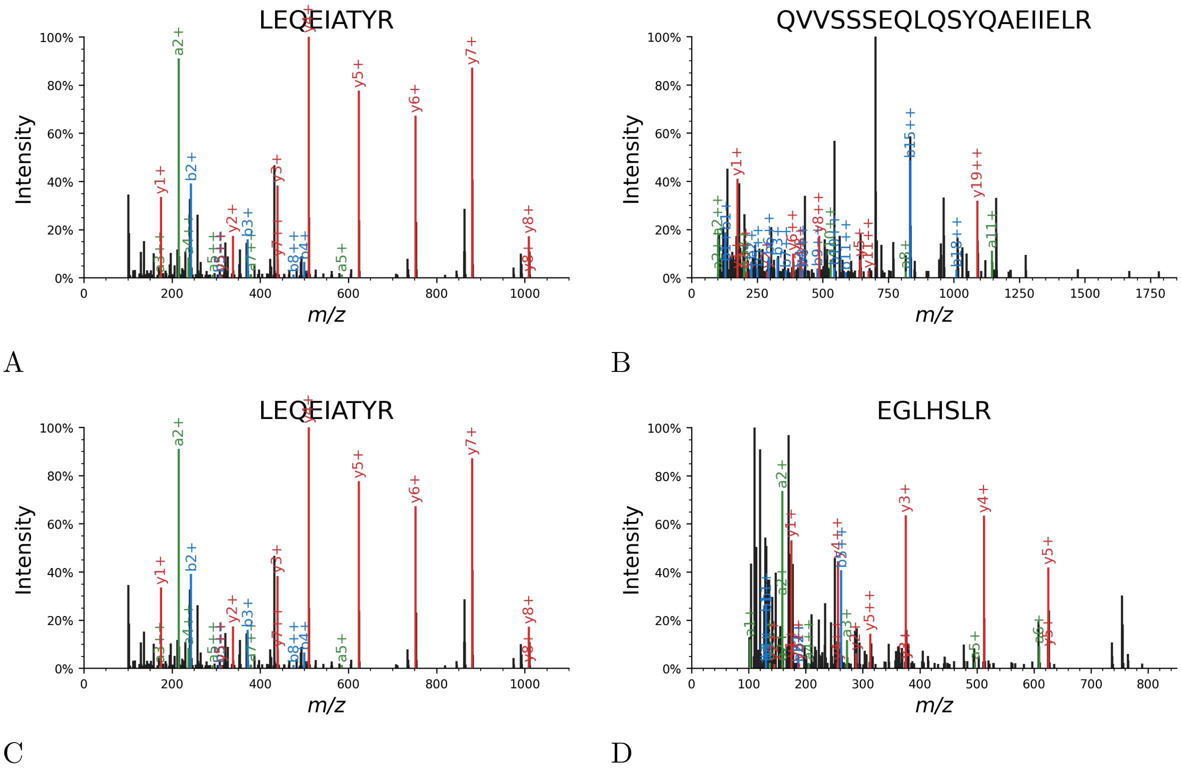


Figure S11: **Annotated spectra for project PXD016001.** Annotated spectra of the best (A) and worst (B) scoring confident detection when controlling the PSM-level FDR using the search-then-select protocol. We also the annotated spectra of the best (C) and worst (D) confident detection when controlling the peptide-level FDR.

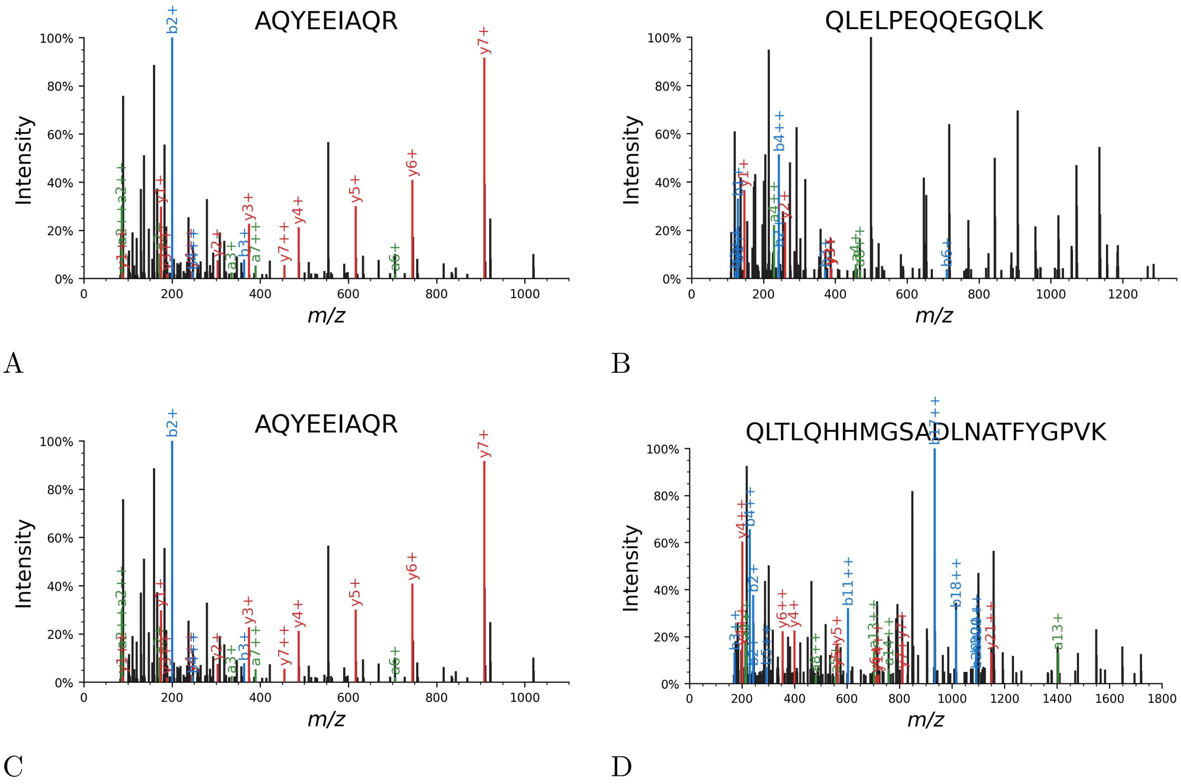


Figure S12: **Annotated spectra for project PXD022965.** Annotated spectra of the best (A) and worst (B) scoring confident detection when controlling the PSM-level FDR using the search-then-select protocol. We also the annotated spectra of the best (C) and worst (D) confident detection when controlling the peptide-level FDR.

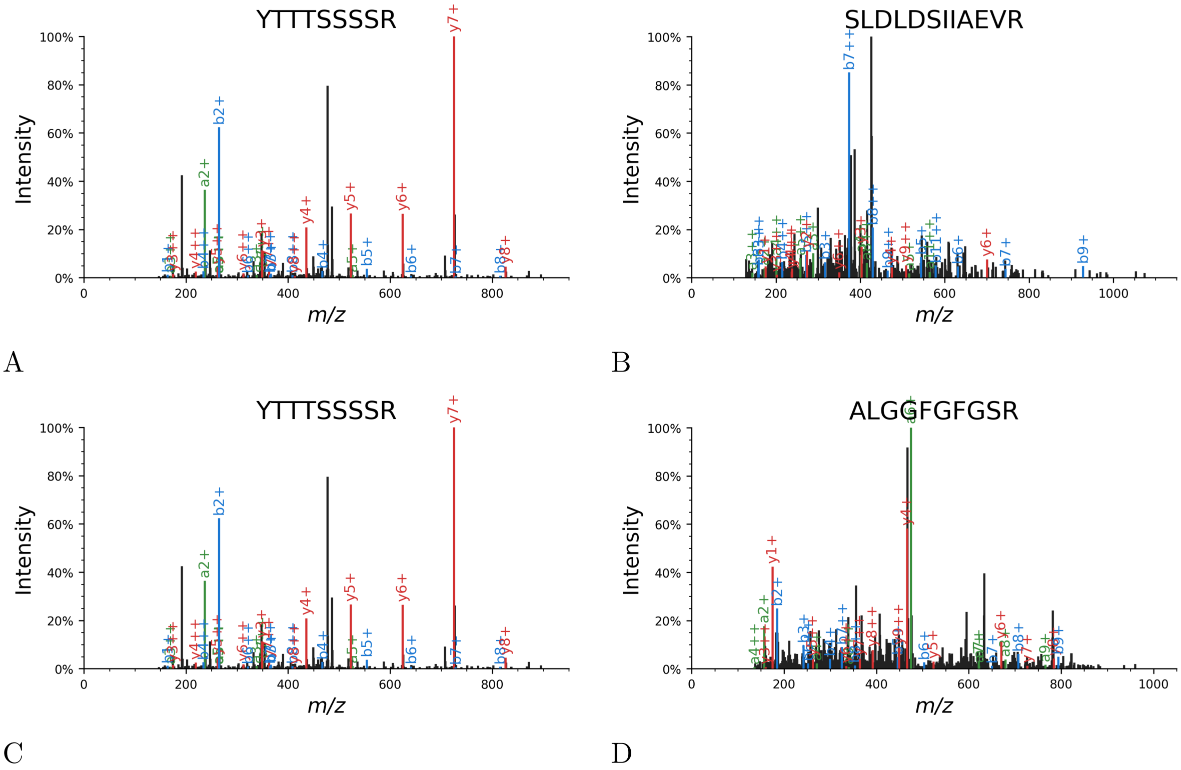


Figure S13: **Annotated spectra for project PXD023729.** Annotated spectra of the best (A) and worst (B) scoring confident detection when controlling the PSM-level FDR using the search-then-select protocol. We also the annotated spectra of the best (C) and worst (D) confident detection when controlling the peptide-level FDR.

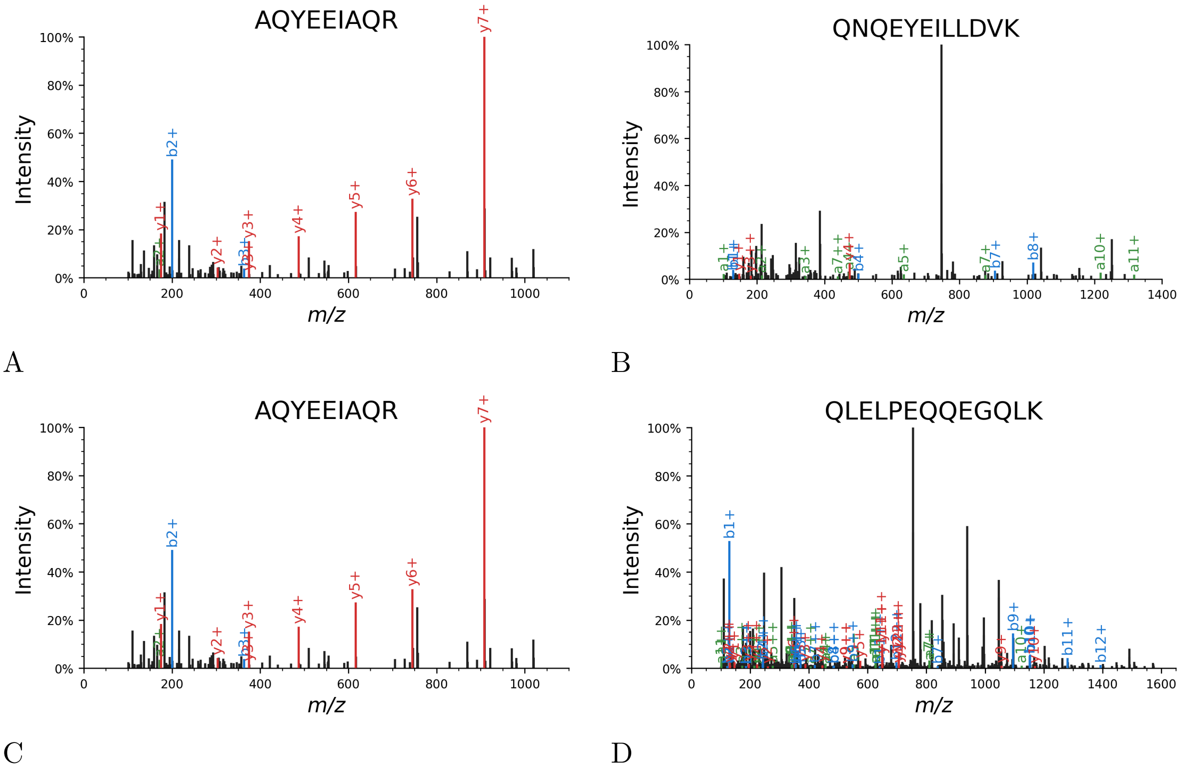


Figure S14: **Annotated spectra for project PXD025088.** Annotated spectra of the best (A) and worst (B) scoring confident detection when controlling the PSM-level FDR using the search-then-select protocol. We also the annotated spectra of the best (C) and worst (D) confident detection when controlling the peptide-level FDR.

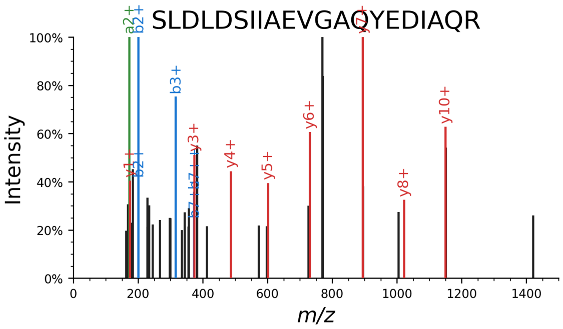


A

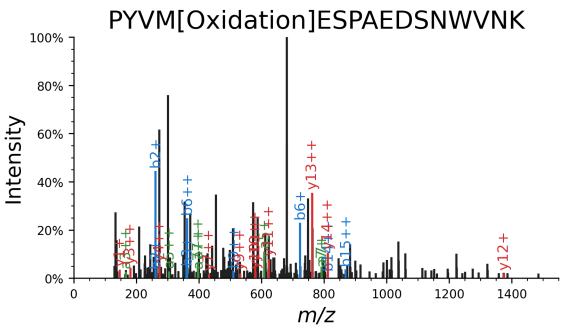


B

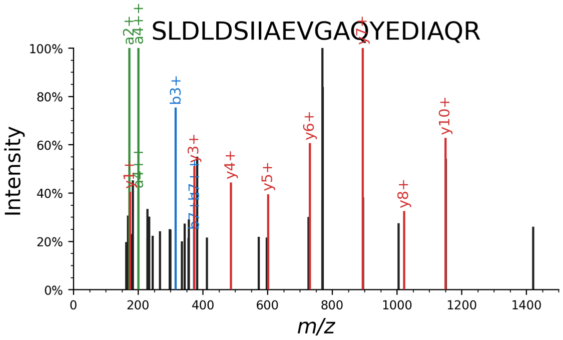


C

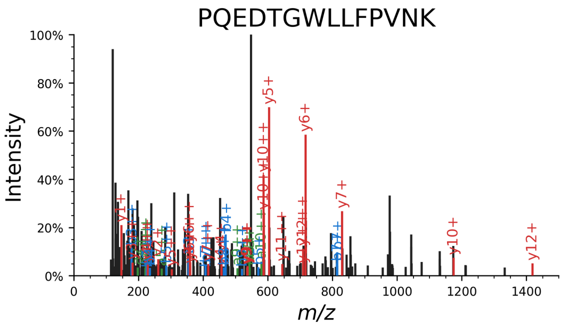


D

Figure S15: **Annotated spectra for project PXD025926.** Annotated spectra of the best (A) and worst (B) scoring confident detection when controlling the PSM-level FDR using the search-then-select protocol. We also the annotated spectra of the best (C) and worst (D) confident detection when controlling the peptide-level FDR.

| PRIDE ID | Species | precursor mass tolerance | score function |
|---|---|---|---|
| PXD010014 | *E. coli* | 28 | XCorr p-value |
| PXD016001 | *E. coli* | 30 | combined p-value |
| PXD022965 | *E. coli* | 32 | combined p-value |
| PXD023729 | *E. coli* | 48 | XCorr p-value |
| PXD025088 | *E. coli* | 22 | combined p-value |
| PXD025926 | *E. coli* | 41 | combined p-value |
| PXD009420 | Yeast | 28 | combined p-value |
| PXD014236 | Yeast | 36 | combined p-value |
| PXD015273 | Yeast | 20 | XCorr p-value |
| PXD015536 | Yeast | 48 | combined p-value |
| PXD015674 | Yeast | 75 | XCorr p-value |
| PXD026806 | Yeast | 34 | combined p-value |

Table S1: **DDA Datasets**. The datasets used in this study that were acquired by DDA.

| PRIDE ID | Species |
|---|---|
| PXD006240 | mouse |
| PXD016668 | mouse |
| PXD019379 | mouse |
| PXD027559 | mouse |
| PXD010126 | *E. coli* |
| PXD012611 | *E. coli* |

Table S2: **DIA Datasets**. The datasets used in this study that were acquired by DIA.

| data acquisition method | sample organism | PRIDE ID | # of runs | # of GVPs |
|---|---|---|---|---|
| DDA (standard search) | *E. coli* | PXD010014 | 37 | 99 |
| | *E. coli* | PXD016001 | 50 | 58 |
| | *E. coli* | PXD022965 | 50 | 36 |
| | *E. coli* | PXD023729 | 14 | 36 |
| | *E. coli* | PXD025088 | 16 | 29 |
| | *E. coli* | PXD025926 | 19 | 21 |
| | yeast | PXD009420 | 16 | 23 |
| | yeast | PXD014236 | 54 | 51 |
| | yeast | PXD015273 | 48 | 74 |
| | yeast | PXD015536 | 135 | 80 |
| | yeast | PXD015674 | 18 | 4 |
| | yeast | PXD026806 | 56 | 41 |
| DDA (subset-neighbor search) | *E. coli* | PXD010014 | 37 | 0 |
| | *E. coli* | PXD016001 | 50 | 8 |
| | *E. coli* | PXD022965 | 50 | 0 |
| | *E. coli* | PXD023729 | 14 | 10 |
| | *E. coli* | PXD025088 | 16 | 5 |
| | *E. coli* | PXD025926 | 19 | 8 |
| | yeast | PXD009420 | 16 | 0 |
| | yeast | PXD014236 | 54 | 0 |
| | yeast | PXD015273 | 48 | 0 |
| | yeast | PXD015536 | 135 | 8 |
| | yeast | PXD015674 | 18 | 0 |
| | yeast | PXD026806 | 56 | 0 |

Table S3: **Number of GVPs per project** A table of the number of detected GVPs per PRIDE project. The false discovery rate was estimated using either the search-then-select or the subset-neighbor-search procedure and filtered to a 5% PSM-level FDR.

| GVP | genes |
|---|---|
| FASFIDK | KRT2, KRT3, KRT4, KRT5, KRT6A, KRT6B, KRT6C, KRT7, KRT8, KRT71, KRT72, KRT73, KRT74, KRT75, KRT76, KRT77, KRT79, KRT84 |
| FLEQQNK | KRT3, KRT5, KRT6A, KRT6C, KRT7, KRT75, KRT76, KRT78, KRT79, KRT8, KRT81 , KRT83, KRT84, KRT85, KRT86, TTBK2 |
| LAADDFR | KRT10, KRT14, KRT15, KRT16, KRT17, KRT19 , KRT24, KRT28, KRT31, KRT32, KRT33B, KRT35, KRT36, KRT37, KRT38 |
| LEQEIATYR | KRT13, KRT15, KRT16, KRT17, KRT19, KRT20 |
| AQYEEIAQR | KRT2, KRT4, KRT6A, KRT6B, KRT6C |
| FLEQQNQVLETK | KRT71, KRT72, KRT73 |

Table S4: **Common GVPs found in other keratin sequences.** A table of GVP sequences and a list of other keratin genes where each sequence is present but is not a GVP.
